# Role of canonical and non-canonical cAMP Sources in CRHR2α-dependent Signaling

**DOI:** 10.1101/2023.10.26.563777

**Authors:** Natalia G. Armando, Paula A. dos Santos Claro, Mariana Fuertes, Eduardo Arzt, Susana Silberstein

**Affiliations:** Instituto de Investigación en Biomedicina de Buenos Aires (IBioBA) - CONICET - Partner Institute of the Max Planck Society, Buenos Aires, Argentina; Universidad Nacional de Quilmes, Departamento de Ciencia y Tecnología, Buenos Aires, Argentina; Departamento de Fisiología y Biología Molecular y Celular, Facultad de Ciencias Exactas y Naturales, Universidad de Buenos Aires, Buenos Aires, Argentina

**Keywords:** CRH system, cAMP sources, CRHR2α, HT22 cells, soluble adenylyl cyclase

## Abstract

Hippocampal neurons exhibit activation of both the conventional transmembrane adenylyl cyclases (tmACs) and the non-canonical soluble adenylyl cyclase (sAC) as sources of cyclic AMP (cAMP). These two cAMP sources play crucial roles in mediating signaling pathways downstream of CRHR1 in neuronal and neuroendocrine contexts. In this study, we investigate the involvement of both cAMP sources in the molecular mechanisms triggered by CRHR2α. Here we provide evidence demonstrating that UCN1 and UCN3 exert a neuritogenic effect on HT22-CRHR2α cells, which is solely dependent on the cAMP pool generated by sAC and PKA activity but independent of ERK1/2 activation. Through the characterization of the effectors implicated in neurite elongation, we found that CREB phosphorylation and c-Fos induction rely on PKA activity and ERK1/2 phosphorylation, underscoring the critical role of signaling pathway regulation. These findings strengthen the concept that localized cAMP microdomains actively participate in the regulation of these signaling processes.

## Introduction

G protein-coupled receptors (GPCRs) constitute the largest protein family within the human proteome, exerting crucial regulatory functions in various cellular processes, including the response to neuromodulators, lipid-based ligands, and peptide hormones, among others. Due to their pivotal involvement in a myriad of physiological pathways, GPCRs have emerged as prime targets for developing novel therapeutic drugs. Traditionally, GPCRs were thought to function via the canonical paradigm, wherein ligand binding triggers the activation of heterotrimeric G proteins, subsequently leading to the activation of transmembrane adenylyl cyclases (tmACs) and a subsequent rise in intracellular cAMP levels (1). However, recent advancements have unveiled the remarkable complexity of GPCR activation mechanisms. These include elucidating G protein-independent signaling pathways and identifying signaling events originating from endosomal compartments (2,3). Thus, our understanding of GPCR signaling has expanded beyond the conventional model, shedding light on the intricate molecular events underlying their activation. cAMP plays a critical role in the intracellular signaling pathways activated by hormones GPCRs. These receptors typically induce an elevation in intracellular cAMP levels, facilitating the activation of protein kinase A (PKA). Subsequently, PKA initiates a cascade of events that culminate in the activation of mitogen-activated protein kinases (MAPKs), with ERK1 and ERK2 (ERK1/2) being the predominant MAPK subgroup involved in brain signaling (4). Moreover, PKA is crucial for phosphorylating the cAMP response element-binding protein (CREB), which serves as a mediator in various transcriptional processes and is vital for integrating on neuronal activity-induced gene programs (5,6). CREB regulates numerous target genes, among which c-Fos holds particular significance in neurons, especially within the hippocampus, due to its crucial role in memory formation (7,8).

The corticotropin-releasing hormone (CRH) family encompasses four neuropeptides: CRH and urocortin 1-3 (UCNs) and two class B GPCRs: CRHR1 and CRHR2. These receptors exhibit differential selectivity for the neuropeptides (9,10). CRH and UCN1 display high affinity for both receptors, whereas UCN2 and UCN3 selectively bind to CRHR2 (9,11).

Two isoforms of CRHR2, alpha (CRHR2α) and beta (CRHR2β) are expressed in rodents, whereas in humans, three isoforms, alpha, beta, and gamma (CRHR2γ), exist. These isoforms differ in their amino-terminal sequence and tissue distribution (12,13). CRHR2α is the predominant isoform in the rodent brain, exhibiting high expression levels in regions such as the olfactory bulb, bed nucleus of the stria terminalis (BNST), lateral septum (LS), ventromedial hypothalamic nucleus, and dorsal raphe nucleus (14–16). On the other hand, CRHR2β is primarily found in peripheral tissues like the placenta, lungs, skin, skeletal muscle, cardiovascular system, and the choroid plexus in the brain (14,17). While CRHR2γ has been identified in human brain tissue, its specific function remains unknown (10,18,19). Regulation of CRH system plays a key role in orchestrating the adaptive neuroendocrine, behavioral, and autonomic responses to stressful stimuli in mammals. On the other hand, CRH elicits pro-anxiogenic responses which are primarily mediated by CRHR1 receptors located in forebrain excitatory circuits (20) although paralel anxiolytic functions are aso mediated by CRHR1 receptors placed in dopaminergic neurons (21). CRH, a peptide synthesized in the hypothalamic paraventricular nucleus, triggers the activation of the hypothalamic-pituitary-adrenal (HPA) axis by stimulating the release of adrenocorticotropic hormone (ACTH) through pituitary receptors (18,22,23). In the canonical view, the UCNs–CRHR2 signaling system is implicated in promoting stress recovery to maintain homeostasis (24,25). However, the precise underlying mechanisms governing these responses remain unclear.

In recent years, our knowledge of CRH-controlled circuits has increased significantly. However, the molecular mechanisms insight CRH-activated receptors have not been parallel and remains comparatively understudied in general, and even more so for the case of CRHR2.

We have previously reported that the CRHR1-mediated cAMP production involves not only the traditional G protein-dependent tmACs activation but also a unique source of cAMP known as the G protein-independent soluble adenylyl cyclase (sAC). Importantly, our findings demonstrated that the morphological changes observed in a hippocampal cell line overexpressing CRHR1 were only attributed to the cAMP pool generated by sAC (2,26). We want to investigate if the CRHR2α has a similar behaviour in the same cellular context.

In this study, we used the mouse hippocampal cell line HT22 as an effective cellular model to investigate the signaling pathways that are activated by CRHR2α, the primary mRNA splicing isoform found in the mouse brain. To gain further insights, we established stable clones expressing the receptor (HT22-CRHR2α cells) and examined the impact of UCNs stimulation on various downstream molecular components regulated by CRHR2. To monitor these intracellular events, we employed Förster resonance energy transfer (FRET)–based biosensor technology. Our findings demonstrate that UCNs elicit an upsurge in cAMP production and PKA activity within HT22-CRHR2α cells, leading to the phosphorylation of ERK1/2 and CREB, as well as the induction of c-Fos expression. In addition, we found that besides the classical tmAC, sAC is essential for ERK1/2 and CREB phosphorylation, as well as the induction of the c-fos immediate early gene. Furthermore, we explored the potential of UCNs to induce differentiation in HT22-CRHR2α cells. Remarkably, our data show that UCN1 and UCN3 trigger neurite outgrowth in these cells, a process dependent on protein kinase A (PKA) activity but independent of ERK1/2 signaling. Significantly, the function of PKA is specifically facilitated by cAMP generated through sAC, representing a clear divergence from tmAC-mediated mechanisms.

## Results

### Characterization of signaling pathways activated by UCN1 and UCN3 in HT22-CRHR2α cells

In previous studies (2,26,27), we have elucidated the crucial involvement of sAC and PKA in CRHR1 signaling in the widely used HT22 cell line as a representative neuron-like cell model.

Although extensive work has been done to understand the intracellular events that occur after CRHR1 activation, we know comparatively little about CRHR2-dependent signaling. To help fill this gap, we first generated HT22-CRHR2α stable clones exhibiting sustained expression of the CRHR2α (refer to the supplementary materials for detailed information). Neither CRHR1 nor CRHR2 exhibited expression in the parental line HT22, as previously documented by Bonfiglio et al. (2013) (27). This approach, which relies on using cell lines that lack the receptor of interest, has been successfully employed in a multiplicity of signaling studies. This is because it allows for the manipulation of the receptor under study without the presence of endogenous receptors that may act as confounding factors.

Next, to achieve a comprehensive molecular characterization of these HT22-CRHR2α clones, we conducted an analysis of the key molecular players activated in general by GPCRs, such as ERK1/2, CREB, cAMP, PKA, and the immediate early gene c-Fos.

Notably, it is crucial to highlight that CRHR2 activation by both, UCN1 and UCN3 (100nM), triggered a typical early onset phosphorylation of ERK1/2 (Fig 1A). Following stimulation, a distinct acute peak was observed at 6 minutes, followed by a gradual decline lasting for a minimum of 40 minutes (Fig 1A). Using the activation peak as a reference time point, concentration-response curves were constructed for UCN1 and UCN3 (Fig 1B). The EC50 values for ERK1/2 phosphorylation were determined to be 43 nM for UCN1 and 733 M for UCN3, indicating their respective potencies. Based on these findings, a working concentration of 100 nM was chosen for both UCN1 and UCN3 in subsequent experiments.

**Fig 1:**
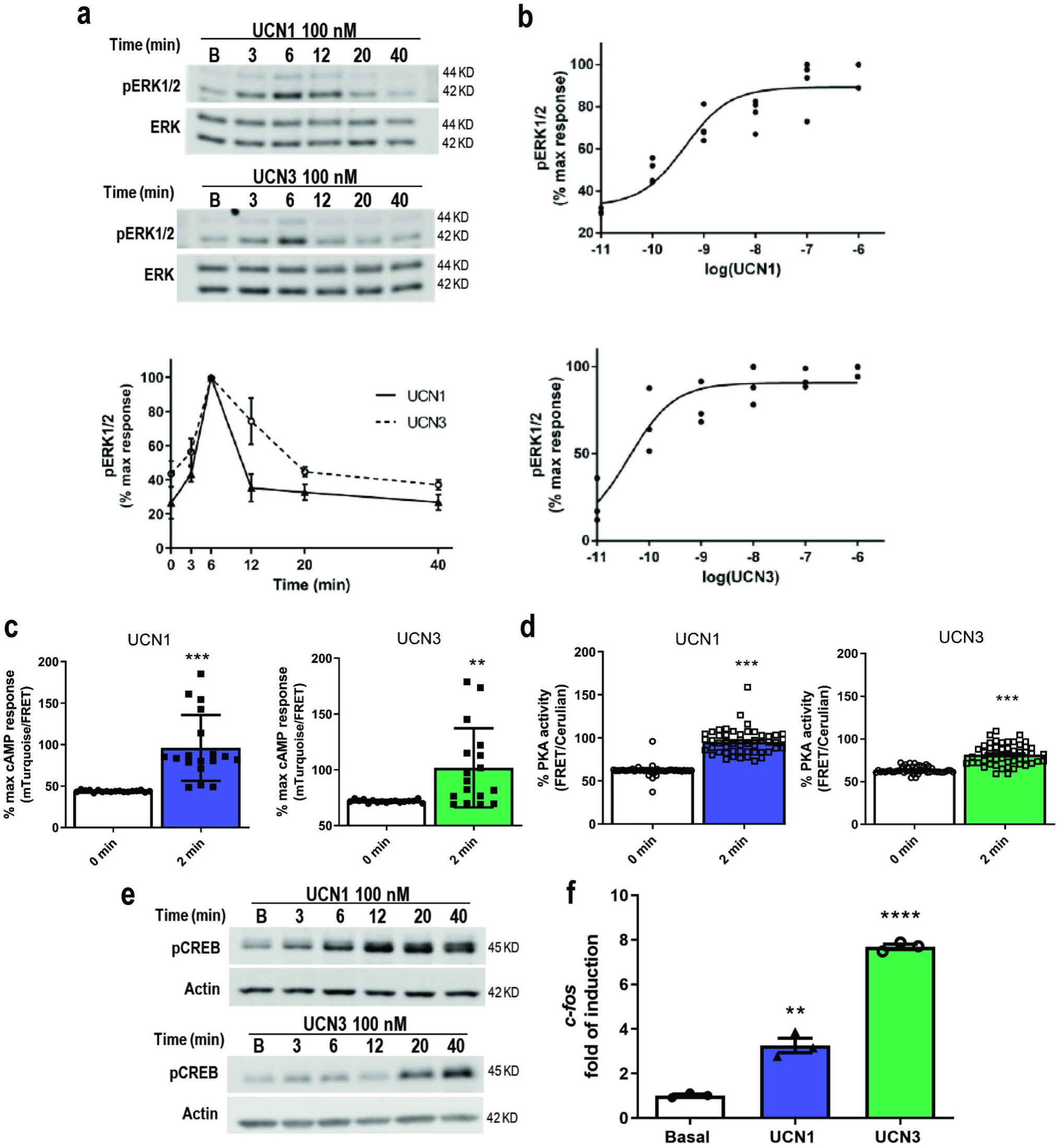
Pathways and signaling effectors activated via CRHR2α using UCN1 and UCN3 in HT22-CRHR2α cells. **a**. Cells were stimulated with 100 nM UCN1 (blue) or UCN3 (green) for the indicated time points. pERK1/2 and total ERK1/2 were determined by western blot and signals were quantified by densitometry as described in Materials and Methods. pERK1/2 was normalized total ERK1/2 for each time point. Results are expressed as the percentage of maximum pERK1/2 obtained at 6 minutes of stimulation. Data: Mean ± SEM from 3 independent experiments. **b**. Concentration-response curves. Cells were stimulated with UCN1 (upper) or UCN3 (bottom) at the indicated concentration for 6 minutes. pERK1/2 and total ERK1/2 were determined by western blot and quantified by densitometry analysis. pERK1/2 was normalized total ERK1/2 in each concentration point. Results are expressed as the percentage of maximum pERK1/2 obtained. Data: Mean ± SEM from 3 independent experiments. To determine EC50 for UCNs-induced ERK1/2 phosphorylation, data were adjusted with 3-parameter dose-response curve. **c-d**. Cells were transfected with FRET biosensors EPAC-S H187 (**c**) or AKAR4 (**d**) and stimulated with 100 nM UCN1 or UCN3. Bars show cAMP response (**c**) and PKA activity (**d**) in basal condition and 2 minutes after stimulation Data: Mean +/- SEM, n=18-30 cells, ** p<0,01 **p<0,001 by student t-test. **e**. Cells were stimulated with UCN1 (top) and UCN3 (bottom) for the indicated time points. pCREB and Actin were determined by western blot. **f**. HT22-CRHR2α cells were stimulated with 100 nm UCN1 or UCN3 for 1 hour. Transcription of c-Fos assesed by real time PCR normalized to HPRT is shown. Data: Mean +/- SEM, n=3, **p<0,01, ****p<0,0001 by t-test.

Our previous studies demonstrated that the activation of ERK1/2 in HT22-CRHR1 cells depends on CRH and requires the presence of cAMP and PKA activity (2). Building upon these findings, we sought to investigate the activation of both cAMP and PKA effectors in HT22-CRHR2α cells using UCN1 and UCN3 as stimuli. To achieve this, we employed FRET-based biosensors, Epac-S^H187^ (a reliable indicator of cAMP levels) and AKAR4 (an indicator of PKA activity) that enable real-time, single-cell measurements (2). FRET signal of Epac-S^H187^ decreases with increasing cAMP concentration, while the FRET signal of AKAR4 increases with elevated PKA activity. To evaluate the response in HT22-CRHR2α cells, we transiently transfected these biosensors and applied UCNs by bath application. Our results revealed a decrease in FRET signal for cAMP response and an increase in FRET signal for PKA activity, providing evidence of heightened cellular cAMP levels (Fig 1C) and augmented PKA activity (Fig 1D).

CREB was initially described as a transcription factor that depends on cAMP response and participates in the maintenance of the long-term effect triggered by the second messengers in the cell (6,28,29). We corroborate the activation of CREB in our system using both urocortins (Fig 1E). In many relevant cell systems, CREB is stalled in promoters of immediate-early genes, ready to rapidly induce transcription (30).

To further explore the functional consequences of increased levels of phosphorylated CREB (pCREB), we examined the induction of its target gene, c-Fos. Real-time PCR analysis revealed a notable increase in c-Fos mRNA expression after 1 hour of UCN1 or UCN3 stimulation (Fig 1F). These findings provide evidence that, downstream of CRHR2α activation in HT22-CRHR2α cells, the stimulation with UCN1 or UCN3 leads to a cascade of events involving cAMP response, PKA activity, ERK1/2, CREB phosphorylation, and ultimately, the induction of the c-Fos gene expression.

#### tmAC and sAC mediate ERK1/2, CREB and c-Fos activation

The transmembrane adenylyl cyclases (tmACs) were the first enzymatic machinery identified as responsible for cAMP production in cells. G proteins regulate these enzymes and can be directly activated by forskolin, a diterpene that has been widely used for this purpose (31,32). It is now accepted the existence of another source of cAMP, sAC, that is insensitive to G protein regulation, but it is directly activated by bicarbonate (33) and calcium ions (34,35). Although sAC was initially described in testis, it is now known to have a ubiquitous expression pattern (36). In our laboratory, we have previously demonstrated high expression of sAC in the mouse hippocampus, cortex, and HT22 cells (2,26). Additionally, it has been reported that sAC and tmACs can tightly regulate distinct downstream pathways from specific microdomains, leading to the spatiotemporal control of internal cAMP pools (3,37–39). Given the observation that cAMP response increases upon UCNs stimulation, we next explored the impact of these two different cAMP sources on the activation of ERK1/2, CREB and the induction of c-Fos mRNA. To evaluate the dependency on tmACs, we introduced the selective inhibitor 2′,5′-dideoxyadenosine (ddA) to the cells. Interestingly, a transient decrease in UCNs-mediated ERK1/2 activation was observed after 5 minutes of stimulation (acute peak), which returned to baseline after 30 minutes (Fig 2 A, B ddA). Remarkably, when the cells were preincubated with the sAC selective inhibitor KH7, a more sustained inhibition was achieved, leading to a reduction in phospho-ERK1/2 levels for up to 30 minutes (Fig 2 A, B KH7). Additionally, the activation of CREB and the induction of c-Fos in response to UCNs stimulation were also impaired in the presence of these inhibitors (Fig 2 C-E). To further support these findings, we conducted experiments using a growth medium lacking bicarbonate, a specific modulator of sAC (S1 Fig), which yielded similar results.

**Fig 2:**
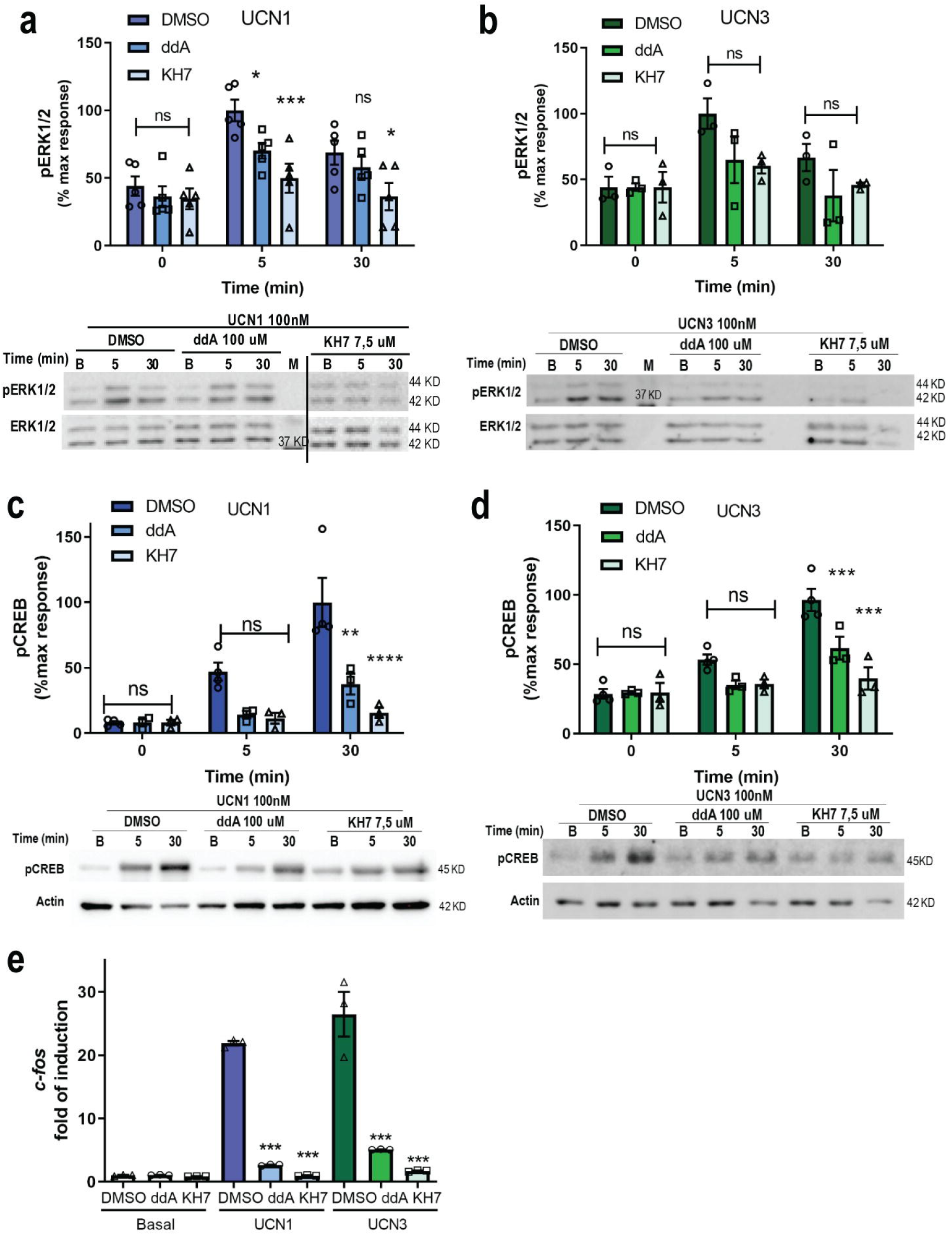
Both cAMP sources are involved in the activation of ERK1/2, CREB and c-Fos. HT22-CRHR2α cells were stimulated with 100 nM UCN1 (blue) or UCN3 (green) for the indicated time points in the presence of tmAC-specific inhibitor (ddA) or sAC-specific inhibitor (KH7) at the indicated concentrations. **a-b**. phosphorylated ERK1/2 and total ERK1/2 and **c-d**, phosphorylated CREB and Actin were measured by immunoblotting and quantified by densitometry using Fiji ImageJ software. pERK1/2 was normalized total to ERK1/2 and CREB to Actin. Results are expressed as the percentage of maximum pERK1/2 or pCREB obtained after stimulation. Data: mean ± SEM, **a**, 5 independent experiments **b-d**, 3 independent experiments,**p<0,01, ***p<0,001, ****p<0,0001 ns: no significative with respect to basal by two ways ANOVA followed by Tukey post test. e, c-fos mRNA levels were determined by RT PCR and normalized to Hprt after 15 minutes of pre-incubation with inhibitors (100 μM ddA or 7,5 μM of KH7) and 45 minutes of stimulation with 100 nM UCN1 or UCN3. Data: Mean ± SEM, n=3,***p<0,001 respect to basal by one way ANOVA following by tukey test.

### PKA and ERK1/2 participate in UCNs-dependent CREB activation

As previously mentioned, intracellular increase of cAMP stimulates CREB, which acts as a crucial effector in neuronal function (40). Similar studies on MAPKs in neurons, reported the involvement of ERK1/2 in CREB activation in PC12 cells stimulated with NGF (41). However, studies by Kuo et al, have indicated that CREB phosphorylation in immortalized hippocampal cells is not influenced by ERK1/2 activation (42). Along this line, we found in previous investigations with HT22-CRHR1 cells that CRH can trigger CREB phosphorylation independently of ERK1/2 activation (26). To explore how this PKA-ERK1/2-CREB cascade behaves downstream CRHR2 in HT22 cells, we conducted experiments pre-incubating cells with the selective inhibitors H89 (PKA) and U0126 (ERK1/2 activation), which resulted in a significant decrease of UCNs-induced CREB phosphorylation (Fig 3 A-D). These findings confirm that both PKA and ERK1/2 play crucial roles in regulating cAMP-dependent CREB activation in response to UCN1 or 3. Also, we corroborate the negative influence of PKA activity blockage in ERK1/2 phosphorylation (S2 Fig). Taken together, these results suggest that the dependency of CREB phosphorylation on ERK1/2 is receptor-dependent even within the CRHRs family.

**Fig 3:**
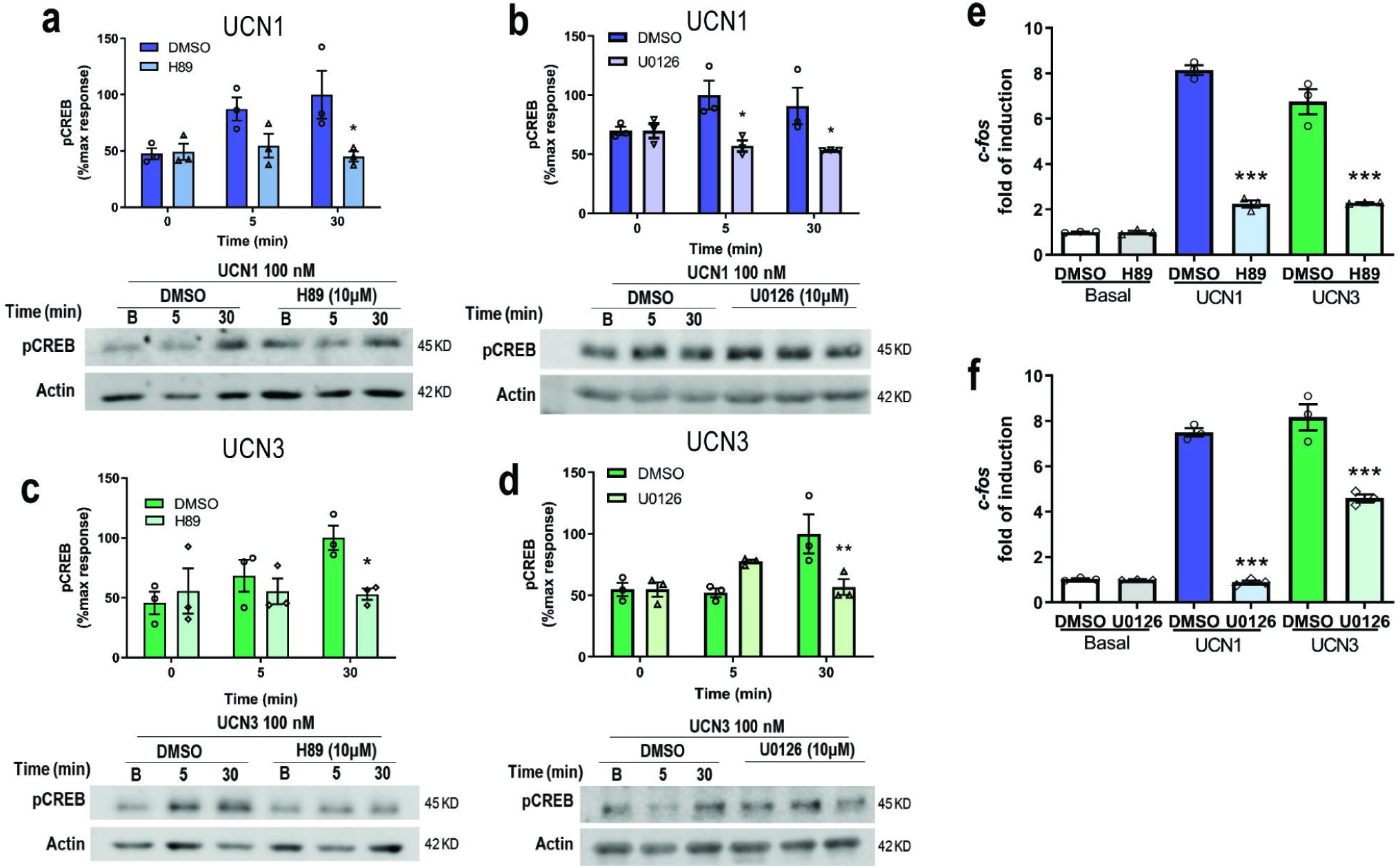
PKA and ERK1/2 activation are critical for CREB phosphorylation in HT22-CRHR2α cells. Cells were stimulated with 100 nM UCN1 (blue) or UCN3 (green) for the indicated time points in the presence or absense of **a, c, e** PKA activity inhibitor (10 μM H89) or **b, d, f** MEK inhibitor (10 μM U0126) **a-d**, CREB and Actin were measured by immunoblotting and quantified by densitometry using Fiji ImageJ software. pCREB was normalized to Actin. Results are expressed as the percentage of maximum pCREB obtained after stimulation. Data: mean± SEM, 3 independent experiments, *p<0,05, **p<0,01 with respect to basal by two ways ANOVA following by Tukey test. **e, f,** c-fos mRNA levels after pre-incubation with inhibitors (10 μM H89 or U0126) and 45 minutes of stimulation with 100 nM UCN1 or UCN3 were determined by RT PCR and normalized to Hprt. Data: Mean ± SEM, n=3, ***p<0,001 with respect to basal by one way ANOVA following by tukey test.

Following the canonical signaling pathway, we next assessed the PKA and ERK1/2 participation in the UCNs-mediated induction of c-Fos in HT22-CRHR2α cells. Our findings demonstrate that the selective inhibitors targeting either PKA or ERK1/2 effectively hindered the UCN1/3-induced expression of c-Fos (Fig 3 E-F). These results provide evidence supporting the crucial role of PKA and ERK1/2 in mediating the UCN-induced c-Fos expression in HT22-CRHR2α cells.

### CRHR2α activation promotes neurite elongation in a sAC- (but not tmAC-) dependent manner

HT22 cells exhibit a characteristic flattened, spindle-shaped morphology, which was also observed in cells expressing CRHR2α. Previous findings from our laboratory documented the morphological changes induced by CRH stimulation in HT22-CRHR1 (26). Intriguingly, we observed similar changes in HT22-CRHR2α cells upon stimulation with UCN1 or UCN3, which led to neurite elongation and the development of more rounded somas (Fig 4A). To quantify these changes, we calculated the differentiation index as the ratio between the length of the longest neurite and the soma diameter for each cell (26). Figure 4A show that a one-hour incubation with UCN1 and UCN3 is sufficient to induce morphological changes indicative of cellular differentiation, mirroring the effects of CRH on HT22-CRHR1 expressing clones. Notably, the HT22 parental line lacks CRHR1 receptor expression, strongly suggesting that the observed effects of UCNs on HT22-CRHR2α cells are a result of genuine CRHR2α activation. Conversely, UCN2 failed to elicit neurite outgrowth in these cells (S3 Fig), implying that this process is ligand-dependent, at least within the context of HT22 cells.

**Fig 4:**
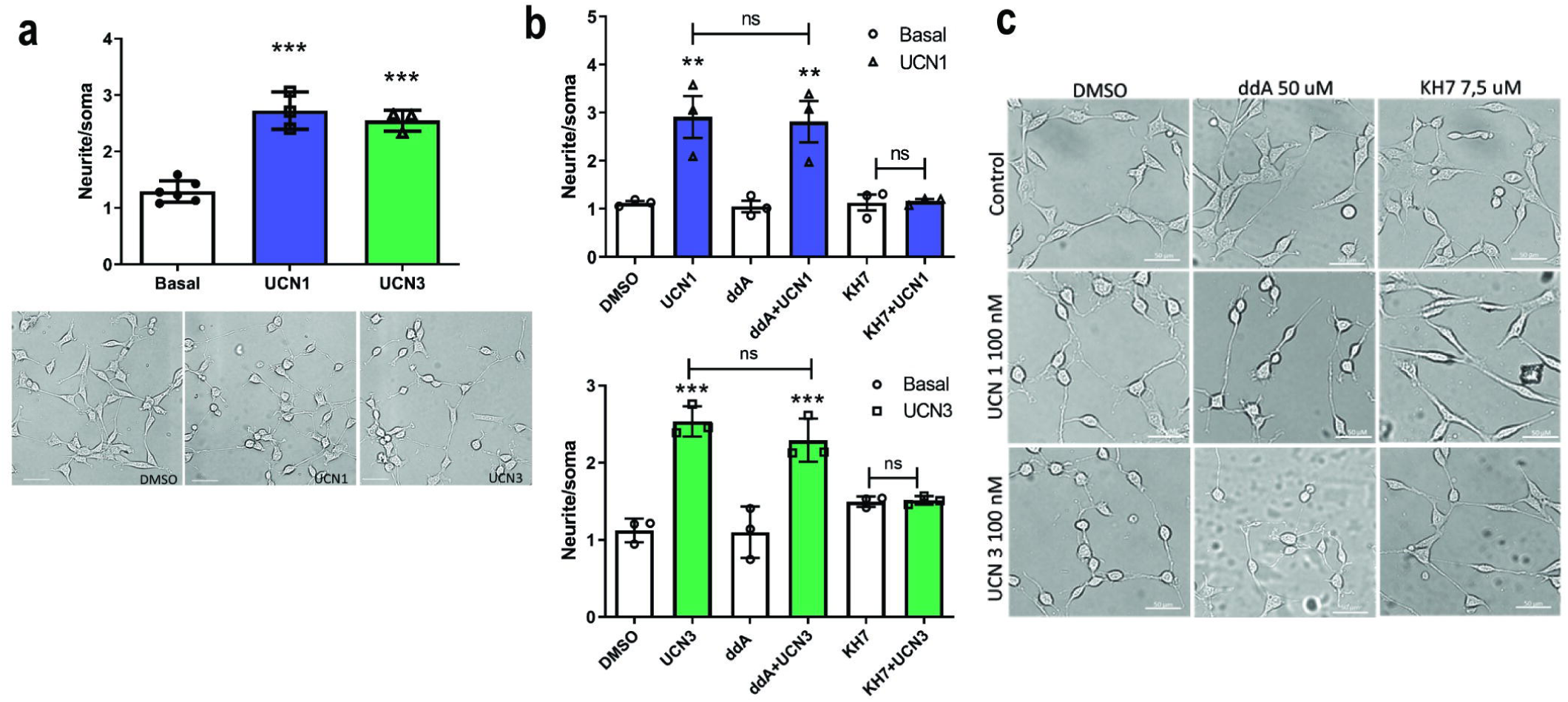
UCNs1/3-induced differentiation of HT22-CRHR2α is SAC-dependent. **a**, HT22-CRHR2α cells were stimulated with 100 nM UCN1 (blue) or UCN3 (green) for 1h. Neurite outgrowth was determined in cells stimulated or in basal conditions. Data: mean± SEM 3 independent experiments, ***p<0,001. Representative photographs are shown for the different treatments. Bars 50 μm. **c-d**, Neurite outgrowth was determined in HT22-CRHR2α cells stimulated with 100 nM of UCN1 **(b**, top) or UCN3 (**b**, bottom) for 1 h in presence of DMSO (control), tmACs-specific inhibitor (50μM 2,5 ddA) and sAC-specific inhibitor (7,5 μM KH7). Representative photographs are shown for each treatment (**c**). Data: mean ± SEM, n = 3. ***, p<0,001 with respect to basal by repeated measures one-way ANOVA following by Tukey test. Bars: 50μM

Previous studies have proposed the involvement of cAMP in GPCR-mediated neurite elongation (43,44). Our previous research demonstrated the essential role of cAMP generated by sAC in the morphological changes induced by CRH in HT22-CRHR1 cells (26). In light of these findings, we investigated whether the cAMP-dependent pathway is involved in the UCN1/3-induced morphological changes observed in HT22-CRHR2α cells. To examine this, we employed specific inhibitors targeting tmAC (ddA) and sAC (KH7) during the stimulation. Surprisingly, pre-incubation with ddA, the tmAC-specific inhibitor, had no effect on the UCNs-dependent morphological changes (Fig 4 B,C). However, when the sAC-specific inhibitor KH7 was included in the stimulation cocktail, the differentiation process was completely blocked (Fig 4 B,C). These results highlight, once again, the existence of CRHR2-specific functions of cAMP sources and utilization.

### CRHR2α-mediated neurite elongation is dependent on PKA activity but independent on ERK1/2 activation in HT22-CRHR2α cells

In order to further investigate the effectors involved in the process of neurite outgrowth, we evaluated UCNs-induced morphological changes in HT22-CRHR2α cells following pre-incubation with various pharmacological inhibitors. Our findings revealed that the neuritogenesis induced by UCNs was completely suppressed by H89, a specific inhibitor of PKA (Fig 5 A,B). Conversely, when the cells were pre-treated with U0126, a specific inhibitor of MEK, no significant differences were observed compared to the control conditions with vehicle treatment (Fig 5 C,D). These results did align with our previous study on CRH-R1-mediated neurite outgrowth in HT22-CRHR1 cells (26), which demonstrated the essential role of PKA in cell differentiation, while the phosphorylation of ERK1/2 was found to be nonessential.

**Fig 5:**
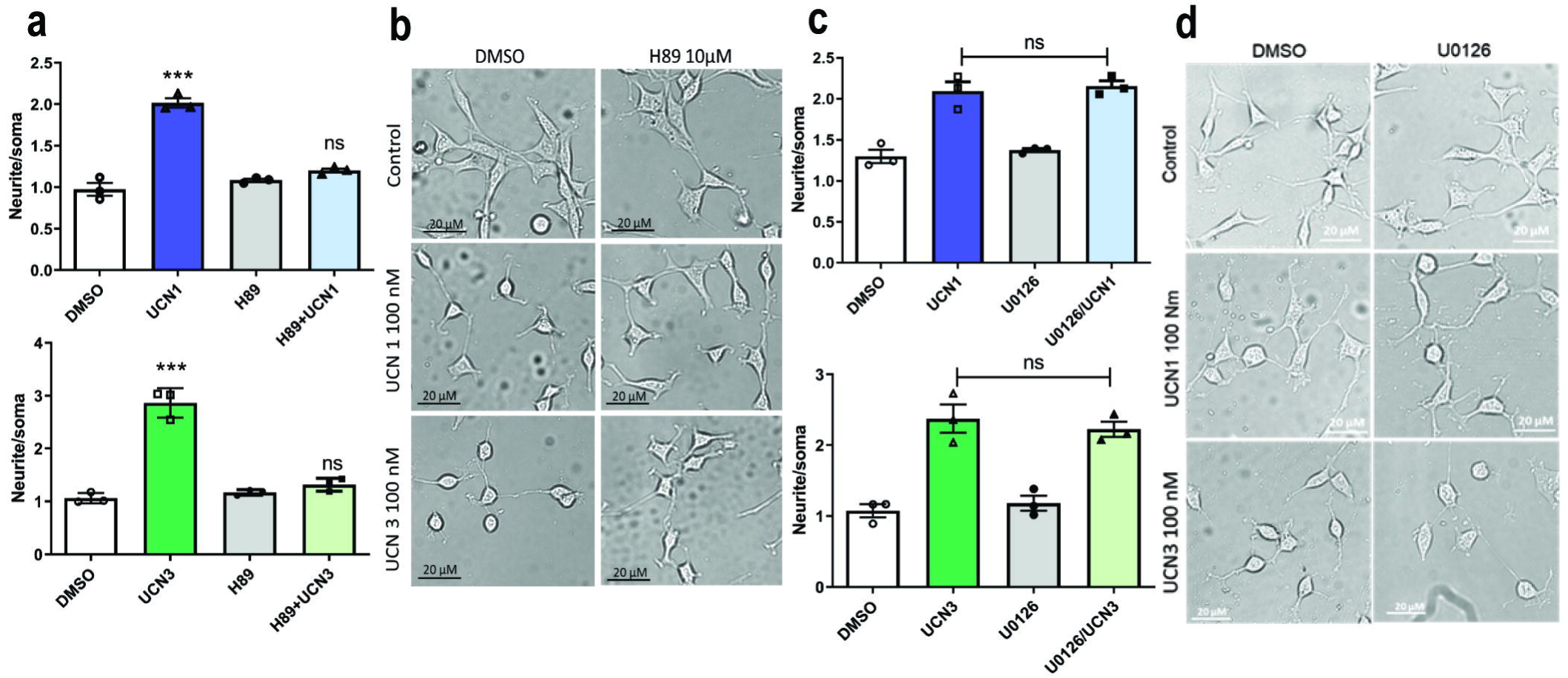
Neurite elongation depends on PKA activity but is independent on ERK1/2 activation in HT22-CRHR2α cells. Cells were stimulated with 100 nM UCN1 (blue) or UCN3 (green) in presence or absence of **a, b**, PKA activity inhibitor (10 μM H89) or **c, d,** MEK inhibitor (10 μM U0126). **a, c**, Neurite outgrowth was determined per cell after 15 min-pretreatment with inhibitors and 1h-treatment with the agonists, as the ratio between the longest neurite and the soma in each cell. Data: mean ± SEM, n = 3. ***, p<0,001 ns: no significative with respect to basal by repeated measures one-way ANOVA following by Tukey test. **b, d**, Representative photographs are shown for each treatment. Bars 20 μM.

Previous studies have demonstrated that activated ERK1/2 is responsible for the differentiation of PC12 cells induced by growth factors (45). However, our findings in HT22-CRHR1 (26) and the present study in HT22-CRHR2α cells offers an alternative view, as we observed ERK1/2-independent cell differentiation. This inconsistency prompted us to investigate whether ERK1/2 activation is always negligible in HT22 cells or solely in the context of CRHR1 (26) or CRHR2 activation. To address this unresolved question, we stimulated HT22-CRHR2α cells with platelet-derived growth factor (PDGF), a known activator of RTK and ERK1/2 in HT22 cells (27). Remarkably, PDGF induced neurite elongation, but unlike UCNs, this effect was completely abolished by pre-incubation with U0126 (S4 Fig), a specific inhibitor of ERK1/2 signaling. These results confirm that the ERK1/2-independence observed in UCN1/3-induced neuritogenesis is a genuine phenomenon.

## Discussion

In this study, our focus was to investigate the signaling pathways of CRHR2α in response to UCNs using hippocampal HT22 stable clones, specifically the HT22-CRHR2α cells and the role of the cAMP sources. Firstly, we discovered that both UCN1 and UCN3 stimulate the phosphorylation of ERK and CREB. Importantly, this phosphorylation event is contingent upon the activation of cAMP and PKA, emphasizing their critical roles as mediators in this process. Furthermore, our results demonstrate that the activation of both ERK and CREB is necessary for the induction of c-Fos, a transcription factor implicated in various cellular processes. Both sources of cAMP were critical for the activation of these effectors. Notably, we also observed that UCN1 and UCN3 promote neurite elongation in HT22-CRHR2α cells. Remarkably, this morphological response is exclusively reliant on the cAMP generated by sAC and is independent of ERK1/2 phosphorylation.

In a previous study by Markovic et al. (2008) (46), it was demonstrated that UCN2 stimulation in HEK293T cells transfected with CRHR2β resulted in the activation of ERK1/2, displaying a similar profile to that observed in HT22-CRHR2α cells upon UCN1 or UCN3 stimulation (Fig 1 A). Considering that CRHR2 receptors belongs to the class B GPCR family as CRHR1, we investigated the involvement of the cAMP canonical pathway by assessing cAMP generation, PKA activity, CREB phosphorylation, and c-Fos induction following UCN1-UCN3 stimulation (Fig 1, 3). Our findings revealed that, not only the canonical tmAC but also the soluble form of ACs regulate the effectors mentioned (Fig 2), similarly to the previously reported effect of CRHR1 in the HT22 cell context (2,26,27). There are limited studies associating sAC with GPCR signaling (47–49). Notably, this is the first report highlighting the involvement of sAC in CRHR2α signaling. Our results suggest that both CRHR1 and CRHR2α activate sAC pathway in the same cellular context. However, this leads us to question their biological significance. We hypothesize that the significance of these receptors lies in their differential trafficking, as previously mentioned. Under basal conditions, the majority of CRHR1 is localized in the plasma membrane (2), whereas CRHR2α is primarily intracellular (50). It is noteworthy that Bonfiglio et al. have delineated a biphasic activation pattern of ERK1/2 in HT22-CRHR1 cells, with each phase governed by different factors. The initial phase hinges upon Braf and PKA, whereas the ensuing phase is reliant on receptor endocytosis and β-arrestins (27). Interestingly, our examination of CRHR2α yielded an absence of a distinct secondary ERK1/2 activation phase (Fig 1 A), which intriguingly contrasts with the phase tied to CRHR1 receptor internalization. This discrepant profile in ERK1/2 activation potentially underpins the divergent localization tendencies of CRH receptors. Notably, this variance in ERK1/2 signaling patterns may substantially account for the observed differences in the subcellular distribution of CRH receptors. This novel insight into the intricate interplay between ERK1/2 signaling profiles and receptor localization provides valuable context for comprehending the dissimilarities in CRH receptor dynamics. No previous studies have explored this aspect, especially considering the ligands involved. It is worth noting that while UCN1 can activate both receptors with similar affinity, CRH and UCN3 are specific to CRHR1 and CRHR2, respectively, particularly at lower concentrations (13,51,52). Further investigations are required to gain a deeper understanding of how these receptors interact within the cell.

c-Fos, an immediate early gene, serves as a widely utilized marker for assessing neuronal activity (53,54). Specifically, within the hippocampus, c-Fos acts as a reliable indicator of activity during behavioural tests associated with memory formation (55–58). Furthermore, stress disrupts memory and learning processes. Although the precise underlying mechanisms remain unclear, evidence suggests a correlation with the functioning of the CRHR1 receptor. Knockout mice lacking this receptor exhibit impaired neuronal plasticity in stressful conditions within the hippocampus (59). On the other hand, Giardino et al. used c-Fos activation to evaluate the function of the CRHR2 in locomotor activity induces by the consumption of methamphetamine in the amygdala (60). Given the interplay between c-Fos, CRHRs, and the activation of CREB transcription factor in HT22-CRHR2α cells derived from the hippocampus, our research was motivated to investigate c-Fos induction within this cellular context. As anticipated, the administration of UCN1 and UCN3 elicited c-Fos expression in HT22-CRHR2α cells (Fig 1F), whereas UCN2 failed to induce this gene expression (S5 Fig). Hale et al. described that UCN2 could generate an increase in c-Fos levels but in serotoninergic neurons from the dorsal raphe nucleus (61). Here we measured c-Fos levels but in a hippocampal cell line, in the hippocampus the 90% of the neurons are glutamatergic. Moreover, we previously demonstrated that sAC is expressed in hippocampal primary cell culture (26) and here we shown that c-Fos induction depends on tmACs and sAC cAMP pools (Fig. 2 E). Thus, we emphasize the importance of further exploration into the actions of c-Fos under CRHRs activation.

We previously described in the HT22 neuronal model the role of the CRHR1 in neuritogenesis (26), here we demonstrated that the CRHR2 has also the same behaviour in the same cellular context (Fig 4A). Since CRHR2 is mostly expressed in specific subsets of neurons in the brain, we propose that the receptor exerts its function on structural plasticity on specific neuronal populations independently of CRHR1. Moreover, not only context is relevant, also the ligand that stimulates the receptor, we observed that UCN1 and 3 evoke this cell differentiation but UCN2 did not elicit similar responses (S3 Fig). Sheng et al. observed a similar behaviour of this urocortin in rat primary hippocampal cell culture, the neuropeptide has an inhibitory effect on the dendrite prolongation (62). On the other hand, Chen et al. demonstrated that UCN2 impairs the action of the Collapsin Response Mediator Protein (CRMP). This protein is expressed during nervous system development and is involved in cellular differentiation (63).

Furthermore, our investigation revealed that PDGF stimulation in HT22 cells leads to neurite elongation, which is contingent upon ERK activation (S4 Fig). It is worth noting that the PACAP receptor, like CRHRs, belongs to the class B GPCR family and activates ERK1/2 through cAMP-PKA signaling without the involvement of sAC (2). On the other hand, PDGFR, a receptor tyrosine kinase (RTK), triggers ERK phosphorylation independently of cAMP (27). These distinctions emphasize the significance of the cAMP pool generated by sAC in the signaling pathways of CRHRs.

Our findings align with recent advances that emphasize the pivotal role of cAMP and its downstream effectors in the regulation of neurite outgrowth. Particularly, in the brain, cAMP-regulated hormone (CRH) emerges as a crucial factor in mediating neurite outgrowth within the noradrenergic locus coeruleus, acting through PKA and ERK1/2-dependent mechanisms (64–66). Conversely, CRH has been shown to impede dendritic arborization in the hippocampus (64,67), highlighting the context-dependent nature of CRHR signaling. The complexities of this system are further underscored by our results and emphasize the importance of selecting the appropriate ligand and cellular context for comprehensive functional investigations of the CRH system, including the differential sources of cAMP that participate in this process.

By considering all these factors, we gain insight into the intricate nature of CRHR signaling and the multifaceted mechanisms underlying neurite outgrowth and dendritic arborization.

Additionally, we provided evidence that both PKA and ERK1/2 activation are essential for CREB activation (Figs 3 A-D). This finding contrasts with the previously reported results for CRHR1 in HT22-CRHR1 cells, where CREB phosphorylation occurs independently of ERK1/2 activation (26). The fact that pERK1/2 does not consistently participate in this process, even when activated in the same cellular context, highlights the specificity of the molecular pathways governing this differentiation process. To gain a comprehensive understanding of the variances observed in CRHRs receptors, it is crucial to continue investigating the intricate mechanisms underlying these divergences.

Our findings in cell differentiation support the idea previously proposed by Swinny and Gounko (65,68) about the capability of the UCNs to evoke morphological changes. Moreover, this is the first report that relates CRHR2α activation with neurite outgrowth in a neuronal cellular context and we report that this depends on the UCNs being used (S3 Fig) and the source where cAMP pool came from.

In summary, the CRHR2α receptor elicits two distinct processes in response to the same stimuli, involving the same actors but operating independently: 1-Activation of tmACs/sAC, leading to PKA-dependent phosphorylation of ERK1/2 and subsequent activation of CREB. 2-Activation of sAC, which is PKA-dependent but ERK1/2-independent, resulting in neurite outgrowth (Fig 6).

**Figure 6:**
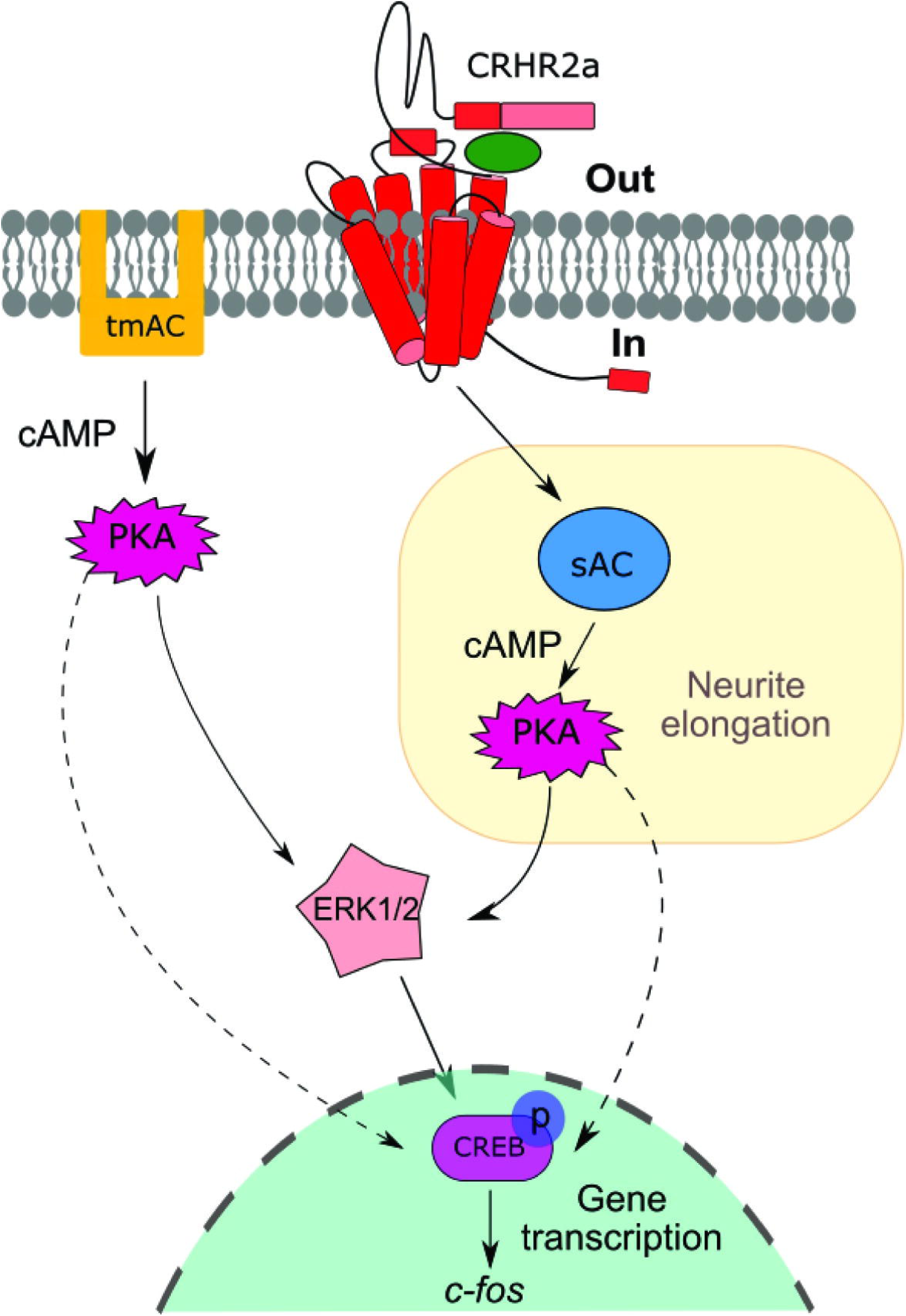
Model proposal of CRHR2α signaling in the HT22 cell context. After stimulating HT22-CRHR2α cells with UCN1 or UCN3, we observed an increase in cAMP levels, leading to the activation of PKA. Additionally, the levels of phosphorylated ERK1/2 and CREB showed a rise, along with the induction of c-Fos. We found that both tmACs and sAC contribute to these effector activations, serving as sources of cAMP. The effectors involved in neuritogenesis in this particular cell context following UCN stimulation are highlighted with a yellow square. The pathways we have confirmed in this study are depicted with bold arrows, while the dashed arrows represent potential pathways that require further investigation.

Given the scarce knowledge on the role of UCNs/CRHR2α in the CNS, the results presented in this work set the ground for the study and characterization of the mechanisms involved in the CRH system activation, which in turn may help understand how its different ligands and receptors work together to mediate stress response.

## Acknowledgments

We thank D. Refojo for his help with the critical reading of this manuscript, A. Attorresi for helping in the use of microscopes and imaging analysis. This work was supported by grants from the Agencia Nacional de Promoción Científica y Tecnológica (ANPCyT, PICT 2017:1977).

## Materials and methods

### Cell culture and generation of stable cell lines expressing CRHR2α

HT22 mouse hippocampal cells were culture as described (22). The lack of tools to track the CRH receptors prompted us to use a construction of the CRHR2α with a FLAG tag on the amino terminal to generate stable HT22-CRHR2α clones. The use of a mouse monoclonal antibody for the FLAG tag allowed us to perform an immunocytochemistry and western blot (data not shown).

The CRHR2α expression vector pcDNA3-FLAG-CRHR2α was kindly provided by Dr Jan Deussing (Max Planck Institute of Psychiatry, Munich, Germany). pcDNA3-FLAG-CRHR2α was used to transfect HT22 cells using Lipofectamine and plus reagent (Thermo fischer scientific) according to the manufacturer’s instructions. For the generation of cell lines stably expressing CRHR2α (HT22-CRHR2α cells), transfected cells were grown in DMEM low glucose (Thermo fischer scientific) in the presence of G418 (800 μg/ml; Life Technologies) to select for transfected cells. G418 surviving clones were subcultured and maintained in DMEM supplemented with 5% (vol/vol) fetal bovine serum (FBS), 4mM L-glutamine (Sigma) 10 mM HEPES (Sigma), 2.2 mg/ml NaHCO_3_, 100 U/ml penicillin (Life Technologies), 100 mg/ml streptomycin (Life Technologies), and 200 μg/ml G418 (Life Technologies) at 37°C in a humidified 5% CO2 incubator. CRHR2α expression levels in the stably transfected cell lines was assessed by Western blot using a mouse monoclonal anti FLAG -antibody (1:5000 dilution; SIGMA). All the experiments were performed at least 3 times with 3 individual clones (clones 4, 13 and 16) with different mRNA expression levels of CRHR2α clones. Using variables clones we ensure that experimental conditions were homogeneous. We did not observe morphological differences between the selected clones.

### Ligand stimulation, drugs, and pharmacological inhibitors

Serum-starved cells were stimulated with human/rat UCN1 (H-3722, Bachem), UCN3 (H-5634, Bachem) or PDGF (01–305; Millipore) at the concentrations and time points indicated. After incubations, cells were washed with ice-cold PBS and maintained in ice. When pharmacological inhibitors were used, cells were pre-treated with the drugs or vehicle 15–30 min before stimulation. The following inhibitors were used: H89 (PKA; 371963 Calbiochem), 2′,5′-dideoxyadenosine (tmACs; 288104, Calbiochem), KH7 (sAC; 3834, Tocris), U0126 (MEK1/2; 662005, Calbiochem). For Western blot assays cells were serum-starved for 6 h in OptiMEM before drug pre-treatments or stimulation. For experiments testing bicarbonate dependence, bicarbonate-free DMEM supplemented with 25 mM Hepes was used to prepare media at 0- and 25-mM sodium bicarbonate concentrations. Each condition was adjusted to physiological pH 7.2 with NaOH and HCl and allowed to equilibrate in the incubator for additional 30 min, and then the pH was checked again and adjusted to 7.2. HT22-CRHR2α cells were serum-starved for 6 h in bicarbonate-free DMEM or 25 mM bicarbonate DMEM before stimulation with 100 nM UCN1 or UCN3.

### Protein detection by immunoblotting

Preparation of cellular extracts and immunoblotting. After treatments, cells were washed with ice-cold PBS and lysed in Laemmli sample buffer. Whole-cell lysates were sonicated and heated to 95 °C for 5min. Samples were resolved by SDS-PAGE and transferred onto 0.45mm nitrocellulose membranes (Millipore) for immunoblotting. Membranes were blocked in TBS-Tween 20 (0.05%) containing 5% inactivated milk at room temperature for 1h under shaking and incubated overnight at 4 °C with the primary antibodies. The following antibodies were used: anti–phospho-ERK1/2 (E-4, sc-7383) from Santa Cruz Biotechnology; anti–total-ERK1/2, (9102, Cell Signaling), anti-phospho CREB (06-519, EMD Millipore), anti-total-CREB (9104, Cell Signaling). Signals were detected by HRP-conjugated secondary antibodies and enhanced chemiluminescence (SuperSignal West Dura, Pierce) using a GBOX Chemi XT4 (Syngene) or by fluorescence IRDye700DX and IRDye800CW secondary antibodies (Rockland). Phosphorylation of MAPK and CREB was detected with the Odyssey Fc Imaging System (Li-Cor Biosystems). Phosphorylated proteins were relativized to its total protein level and results expressed as the percentage of maximum of the phosphorylated protein after stimulation. Immunoreactive signals were analysed digitally using Fiji software.

### Neurite outgrowth assay

Cells seeded in a 40% density in glass-bottom 24-well plates were stimulated with 100nM UCN1 or UCN3 in the presence of vehicle or specific inhibitors in OptiMEM (Gibco). After 1h-treatment, cells were imaged under bright field illumination using an Olympus IX81 inverted epi-fluorescence microscope using a 20X air objective and Zeiss Black software for image acquisition. For each treatment, at least 15 random fields were imaged. Morphological changes quantification was performed using Simple Neurite Tracer plugin for FIJI software. Neurite outgrowth was determined as the ratio between the longest neurite and the soma diameter per cell after 1, measuring at least 100 cells per treatment. For statistical analysis, repeated measures one- or two-way ANOVA followed by the indicated post test (n=3) were performed.

### Spectral Förster Resonance Energy Transfer (FRET) live imaging of the cAMP response and PKA activity

HT22-CRHR2α cells were transfected with FRET biosensors and were seeded in glass-bottom dishes. Cell imaging was performed on an inverted Zeiss LSM 710 confocal microscope (Carl Zeiss Microscopy GmbH) and ZEN Black 2011 software as previously described (2). Images were acquired with a 40x/1.2 water immersion and temperature corrected objective lens at 1024×1024, 16-bit, pixel dwell time of 3.15 μs, with open pinhole (600 μm). For FRET experiments, cells were illuminated with a 30mW 405nm diode laser at 2% laser power, a 405nm dichroic mirror was used and the emission was collected between 413–723nm wavelength (460–500 nm for Turquoise and 515–615 for Venus), every 15 s for a duration of 15min. The saturation level was verified for each image. Phenol red–free DMEM medium supplemented with 20 mM HEPES was used and imaging was performed at 37 °C and 5% CO2. Around 2.5 min after the start of the experiment for setting basal levels, UCN1 or UCN3 were added to the final concentration indicated. The cAMP and the PKA activity response is shown as bars, in which the maximum response measured in a 2-min interval is presented. The data is expressed as percentage of the maximum response, being 100% UCNs-elicited cAMP or PKA activity in control conditions.

### RT-PCR and quantitative real-time PCR

Total RNA was extracted from cell lines, TRIzol reagent (Invitrogen) and complementary DNA synthesis was carried out using M-MLV reverse transcriptase in the presence of RNasin RNase inhibitor (Promega). PCR primers are all intron spanning. Quantitative real-time PCR was performed with Taq DNA polymerase (Invitrogen) and SYBR Green I (Roche) using a CFX96 Touch™ Real-Time PCR Detection System. Relative expression was calculated for each gene by the Ct method with *Hprt* for normalization. Sequences and expected product sizes are as follows:

*hCrhr2* (233 bp)
sense 5′-GACGCGGCACTGCTCCACAG-3′,
antisense 5′-GCATTCCGGGTCGTGTTGT -3′
*c-fos* (172 bp)
sense 5′-ATCGGCAGAAGGGGCAAAGTAG-3′,
antisense 5′-GCAACGCAGACTTCTCATCTTCAAG-3′
*Hprt* (139bp)
sense 5′-TGGGCTTACCTCACTGCTTTCC-3′,
antisense 5′-CCTGGTTCATCATCGCTAATCACG-3′

### Statistics

Each experiment was performed at least 3 independent times. The results are presented as the mean±SEM of each measurement. Comparisons between treatments were performed using Student’s t-test, one or two-way ANOVA (GraphPad Prism) followed by post-hoc tests stated in the Figures. Statistically significant differences are indicated.

## Supporting Information

S1 Fig. **sAC participation in ERK1/2 and CREB activation in HT22-CRHR2α.** Cells were stimulated with 100 nM UCN1 (blue) or UCN3 (green) for the indicated time points in the presence or absence of bicarbonate (25 mM HCO_3_^-^) **a-b**, phosphorylated ERK1/2 and total ERK1/2 and **c-d** phosphorylated CREB and Actin were measured by immunoblotting and quantified by densitometry using Fiji ImageJ software. pERK1/2 was normalized total to ERK1/2 and pCREB to Actin. Results are expressed as the percentage of maximum pERK1/2 or pCREB obtained after stimulation. Data: Mean ± SEM, 3 independent experiments, **p<0,01, ***p<0,001 respect to basal by two ways ANOVA following by Tukey test.

S2 Fig. **PKA participates in ERK1/2 phosphorylation in HT22-CRHR2α cells**. Cells were stimulated with 100 nM UCN1 for the indicated time points in the presence or absence of PKA activity inhibitor (10 μM H89), pERK and total ERK1/2 were measured by immunoblotting and quantified by densitometry using Fiji ImageJ software. pERK1/2 was normalized to total ERK. Results are expressed as the percentage of maximum pERK1/2 obtained after stimulation. Data: Mean ± SEM, 3 independent experiments, *p<0,05 respect with to basal by two ways ANOVA following by Tukey test.

S3 Fig. **UCN2 is unable to differentiate HT22-CRHR2α cells**. **a**, HT22-CRHR2α cells were stimulated with 100 nM UCN2 for 1h. Neurite outgrowth was determined in cells stimulated or in basal conditions. Data: Mean ± SEM 3 independent experiments, ns: no significative. **b**, Representative photographs are shown for the different treatments. Bars 50 μm.

S4 Fig. **Cell differentiation mediated by PDGF in HT22-CRHR2α cells.** Cells were stimulated with PDGF (10 ng/ml) in the presence or absence of MEK inhibitor (10 μM U0126). **a**, Neurite outgrowth was determined per cell after 15 min-pretreatment with inhibitors and 1h-treatment with the agonists, as the ratio between the longest neurite and the soma in each cell. **b**, Representative photographs are shown for each treatment. Data: Mean ± SEM, n=84-106 cells.

S5 Fig. **The induction of c-Fos expression remains undetectable in the presence of UCN2.** HT22-CRHR2α cells were stimulated with UCN2 100 nM ligands concentration for 1 hour. Transcription of c-Fos assesed by real time PCR normalized to HPRT is shown. Data: Mean ± SEM, 3 independent experiments, ns= no significative with respect to basal by two ways ANOVA following by Tukey test.

## Notes

### Competing Interest Statement

The authors have declared no competing interest.

