## Supplementary figures and images for "Role of canonical and non-canonical cAMP Sources in CRHR2α-dependent Signaling"

### Supplemental Figure 1

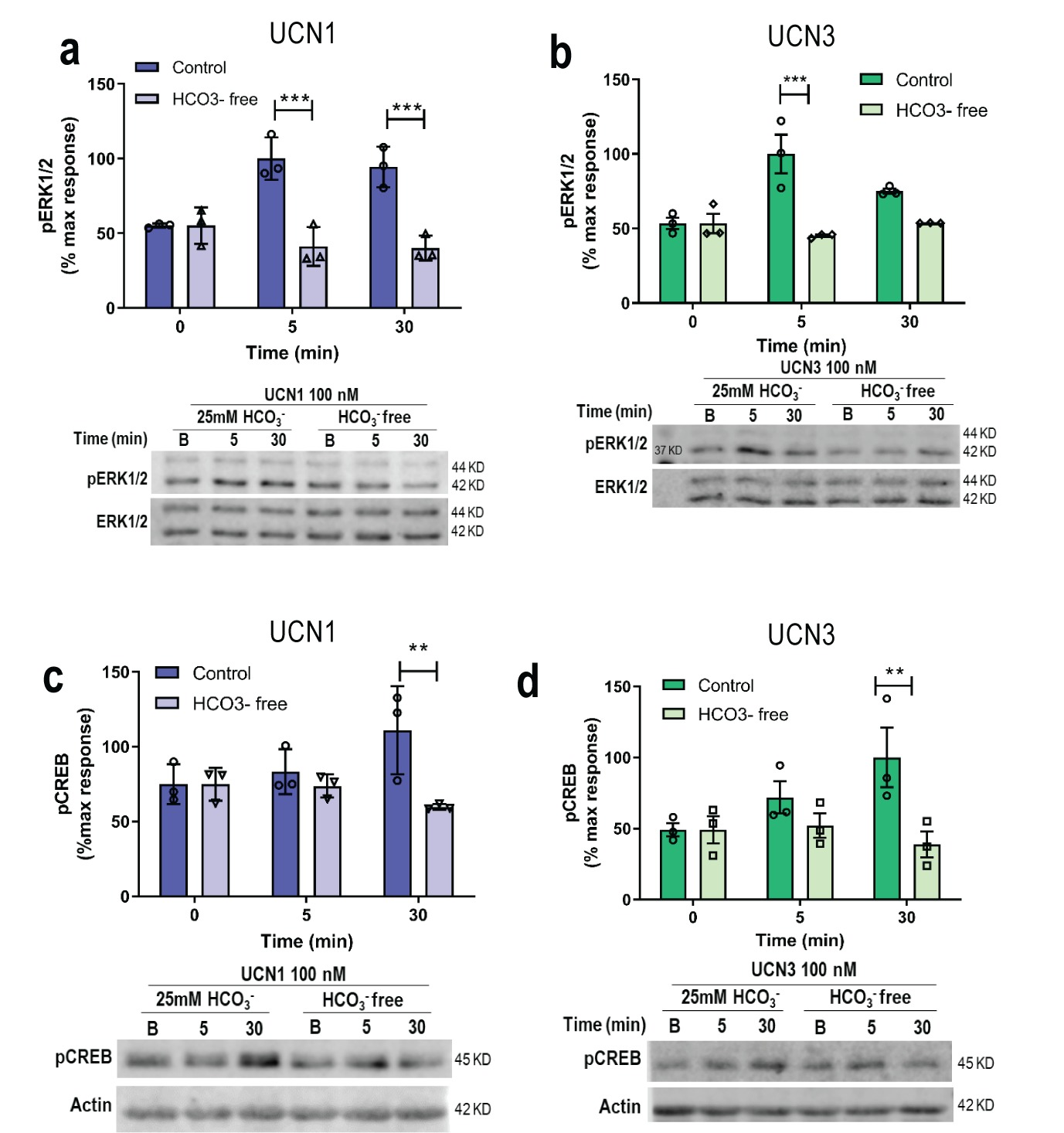

### Supplemental Figure 2

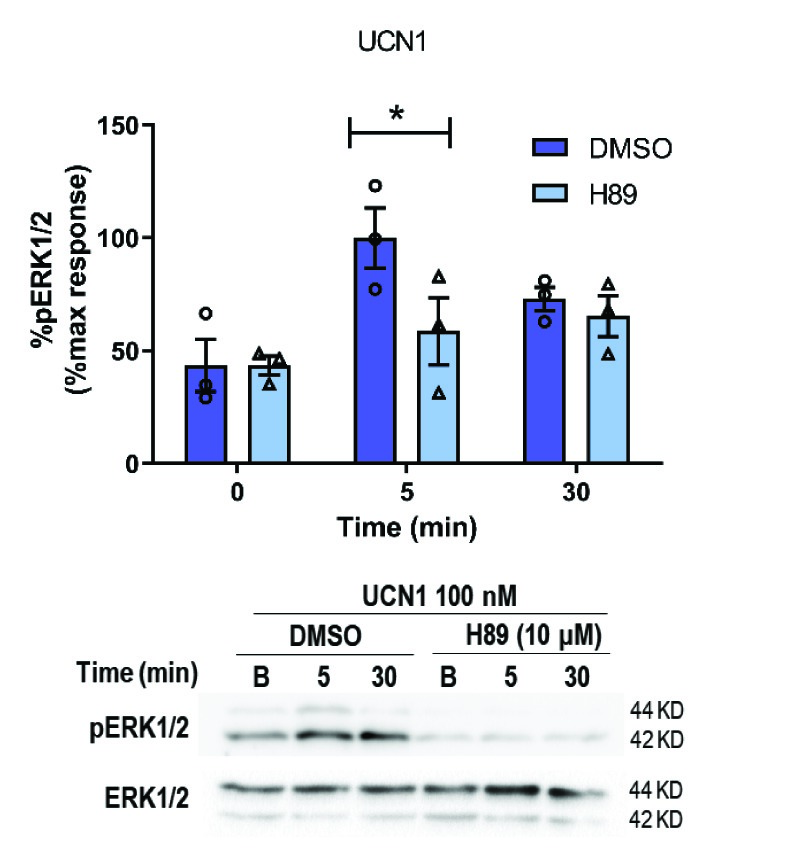

### Supplemental Figure 3

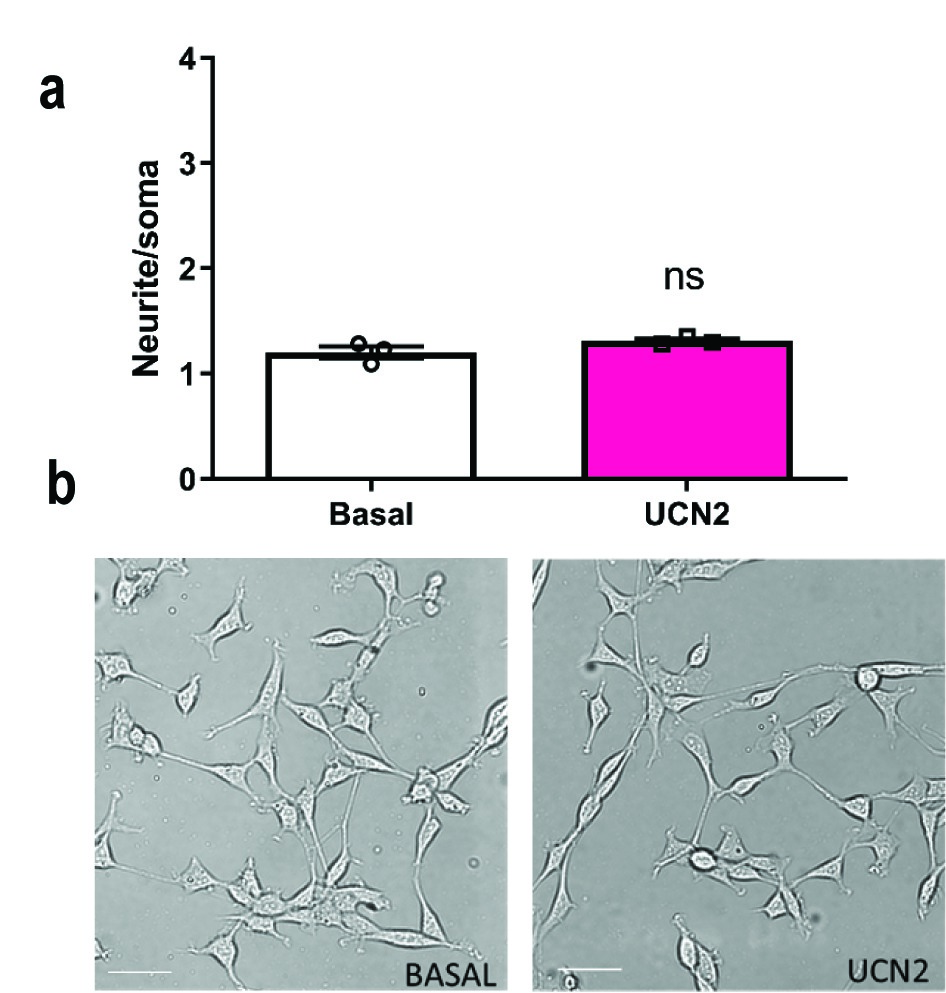

### Supplemental Figure 4

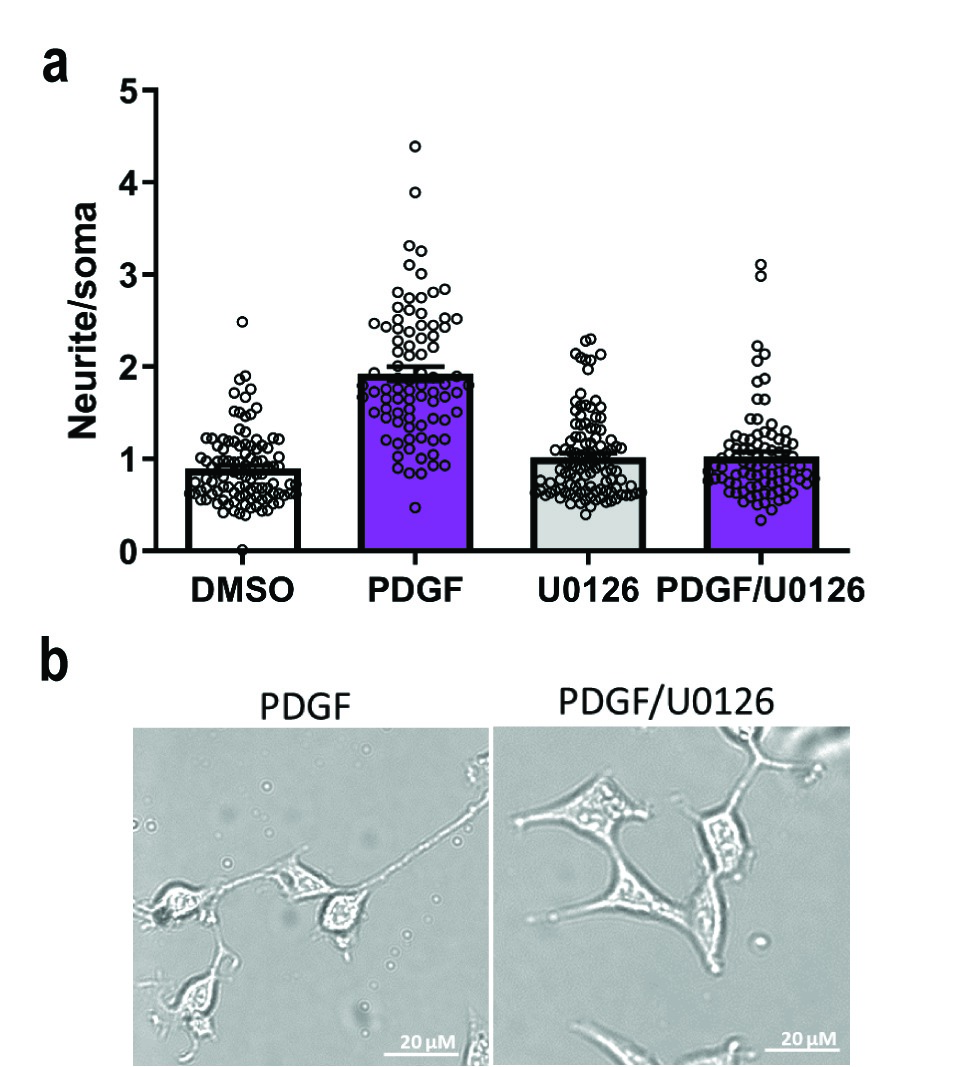

### Supplemental Figure 5

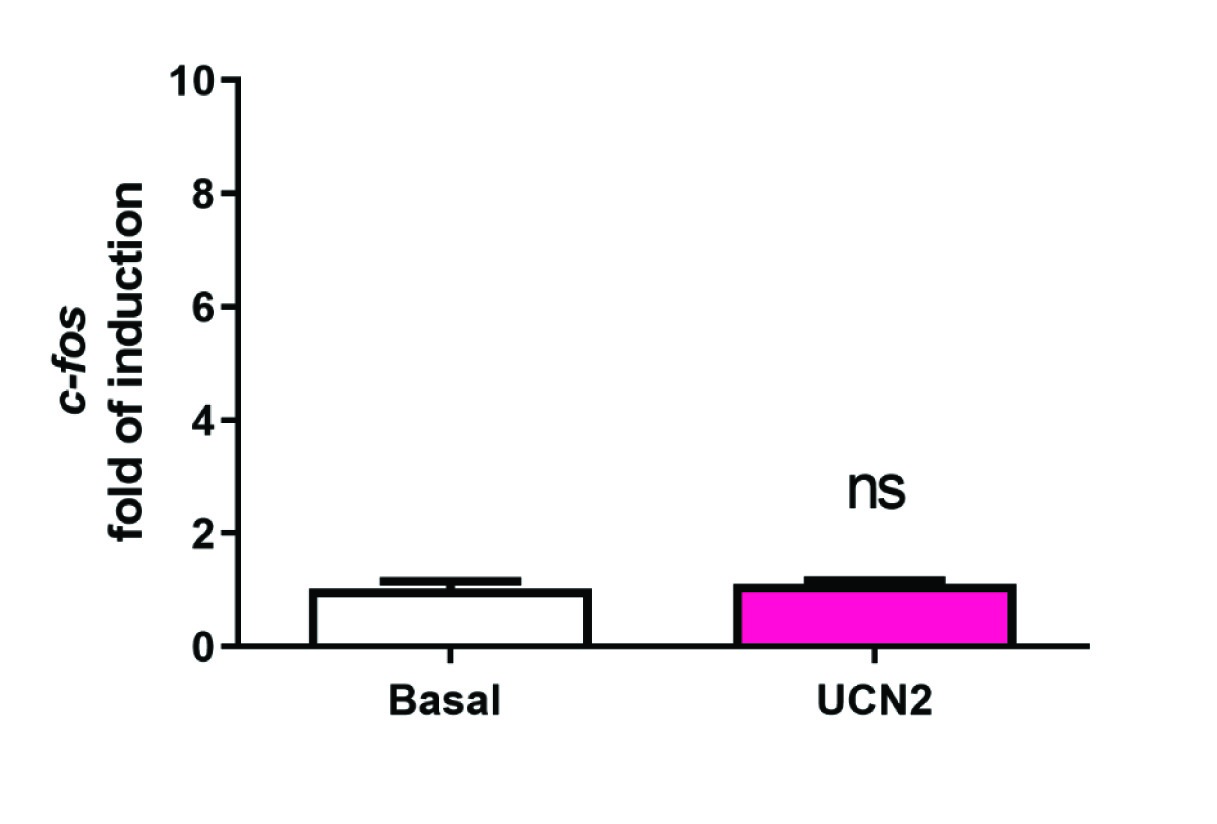
